# Population genomics of a biocontrol agent: Insights into post-introduction establishment in parthenium beetle

**DOI:** 10.64898/2026.06.02.729737

**Authors:** Ranjit Kumar Sahoo, Karthikeyan Vasudevan

## Abstract

Biological control agents often experience demographic bottlenecks during introduction, which can reshape genetic diversity and inbreeding pattern influencing establishment and long-term ecological success in the introduced populations. The leaf-feeding beetle *Calligrapha* (*Zygogramma*) *bicolorata*, introduced in multiple countries across the globe to control the weed *Parthenium hysterophorus*, provides an opportunity to examine how introduction bottleneck and post-introduction breeding practices shape genomic variation and inbreeding patterns. We analyzed whole-genome variation in the introduced beetle population in India by sampling six regions encompassing its current distribution in the country. Using genome-wide variation data, we assessed population structure, genetic diversity, and inbreeding patterns across regions, and inferred historical changes in effective population size to reconstruct post-introduction demographic trajectories. The analyses reveal subtle genetic structure across regions, with overall genetic diversity relatively low compared to other invasive and biocontrol insects. Inbreeding patterns vary among populations, with some regions exhibiting higher cumulative runs of homozygosity than others. Notably, regions subjected to intensive propagation of beetle populations show elevated signature of inbreeding alongside reduced historical effective population size. These results underscore the dynamic genomic consequences of biocontrol introduction and subsequent breeding practices, providing insight into the evolutionary trajectories of introduced biocontrol agents.

## Introduction

To mitigate the impacts of non-native invasive species, biocontrol strategies rely on the introduction of natural enemies sourced from the invader’s native range (Thomas and Willis 1998; Thomas and Reid, 2007; Seastedt 2015). These approaches are generally favored over chemical and mechanical methods because of their economic efficiency and ecological compatibility (Schaffner et al., 2020; van Wilgen et al., 2020). The latter is largely driven by the specificity of ecological interactions between the biocontrol agent (BCA) and the target species, typically manifested through plant–herbivore, predator–prey, or host–parasite relationships (Galli et al., 2024; Stenberg et al., 2021). To reduce unintended ecological consequences, BCAs are often selected for a high degree of specialization (Heimpel and Cock, 2018; Brodeur, 2012). While such introductions have successfully suppressed target populations in many cases, there are documented instances where non-target impacts have raised concerns regarding the long-term ecological sustainability of biocontrol interventions (Louda et al. 1997; Thomas and Reid, 2007; De Clercq et al., 2011; Driesche and Hoddle 2016; He et al. 2021). Therefore, host specificity has become a central focus in biocontrol research, as it is widely assumed to underpin successful outcomes (Heimpel and Cock, 2018; Hoelmer and Kirk, 2005). However, the relationship between specificity and effectiveness remains unresolved. For instance, theoretical models indicate that specialist BCAs may only achieve effective control in target populations exhibiting strong Allee effects (Fagan et al., 2002), implying that outcomes can be highly species-dependent. Despite these insights, the mechanisms through which genetic and demographic factors jointly influence the establishment and spread of BCAs are still not fully elucidated.

Introduced populations of BCAs are typically founded by a limited number of individuals, which are subsequently mass-reared to achieve sufficient numbers for host-specificity testing and eventual field release. Such founding conditions, coupled with inbreeding during propagation, are likely to reduce genetic diversity, thereby constraining adaptive potential and increasing extinction risk. Despite these theoretical expectations, many BCAs have successfully established and proliferated in their introduced ranges (Schwarzlander et al., 2018), presenting a genetic paradox. Originally proposed in the context of invasive species, this paradox refers to the persistence and expansion of populations despite reduced genetic variation at the time of introduction (Estoup et al., 2016; Schrieber and Lachmuth, 2017). Empirical studies across diverse invasive taxa and BCAs have questioned the generality of this concept (Schrieber and Lachmuth, 2017; Hufbauer and Roderick, 2005). Nonetheless, a recurring pattern suggests that factors such as high propagule pressure, repeated introductions, population admixture, or combinations thereof can mitigate the loss of genetic diversity in founding populations. Evidence from experimental and modelling studies in BCAs further underscores the importance of propagule pressure in facilitating establishment success (Fauvergue et al., 2012). However, in practical biocontrol programs, stringent regulations governing transboundary movement of organisms – such as Convention on Biological Diversity and Nagoya protocol (Cock et al., 2010; Deplazes-Zemp et al., 2018) – along with mortality during transport, often constrain the effective propagule size. Hence, the relative contributions of demographic processes and genetic factors in shaping genetic diversity during BCA introduction, establishment, and subsequent expansion remain insufficiently resolved.

In this study, we employ the Mexican leaf beetle, *Calligrapha* (*Zygogramma*) *bicolorata* Pallister, 1953 (Coleoptera: Chrysomelidae) (Shawn et al., 2024), as a model system for biocontrol genomics. Native to tropical America, this species has been deliberately introduced into multiple regions across Asia, Africa, and Australia for the management of the invasive weed *Parthenium hysterophorus* (Dhileepan and Strathie 2009; Dhileepan and Wilmot Senaratne 2009; CABI 2021). *C. bicolorata* exhibits several attributes desirable in an effective biocontrol agent, including a short generation time, high fecundity, and strong host specificity (Jayanth, 1991). Notably, it is among the few insect BCAs of weed species for which a fully assembled reference genome is available (Sahoo et al., 2003), making it particularly suitable for advancing genomic investigations in biocontrol systems.

Here, we focus on populations established in India, where the species was first introduced in 1984 through a single consignment of approximately 66 live individuals (out of total 307) imported from Mexico (Sushilkumar, 2009; Jayanth, 1987). Since then, its distribution has expanded substantially, aided by mass-rearing programs across multiple breeding centers and subsequent augmentative field releases (Mishra, 2012; Sushilkumar, 2009; Jayanth and Nagarkatti, 1987). While such practices likely facilitate rapid geographic spread and enhance gene flow among populations, they may also intensify inbreeding effects due to repeated propagation from limited stock. To investigate these dynamics, we performed whole-genome resequencing of 74 individuals sampled from six regions across the country. This dataset was used to assess contemporary patterns of genetic diversity and population structure at the genome-wide level, as well as to infer recent demographic history. Specifically, we evaluate how the initial introduction bottleneck, together with recent breeding and augmentation practices, has shaped present-day genetic diversity in this species.

## Results

### Variant calling

We generated whole-genome resequencing data from 74 individuals sampled across the distribution of *C. bicolorata* in India, spanning six distinct regions (KA, OD, WB, RJ, CH, and MP) (Fig. 1A; details in Table S1). Of these, 24 individuals (four per region) were sequenced at high depth, yielding an average of 30.9 Gb of paired-end reads per individual. The remaining 50 individuals (with at least eight representatives per region) were sequenced at lower depth, with an average of 5.6 Gb per individual. Trimmed reads were mapped independently to the *C. bicolorata* reference genome (GCA_032362365) (Sahoo et al., 2023). This resulted in a mean coverage of 27.6X (93% breadth) for the high-coverage dataset and 4.9X (87% breadth) for the medium-coverage dataset. The high-coverage dataset was downsampled to 6X to facilitate joint variant call of all 74 samples. Variants located within annotated repeat regions, sex-linked contigs, and contigs shorter than 100 kb were excluded. Based on the kinship analysis, nine individuals were removed from the dataset. The final dataset comprised 65 individuals and 39,95,788 high-confidence biallelic SNPs.

**Figure 1.**
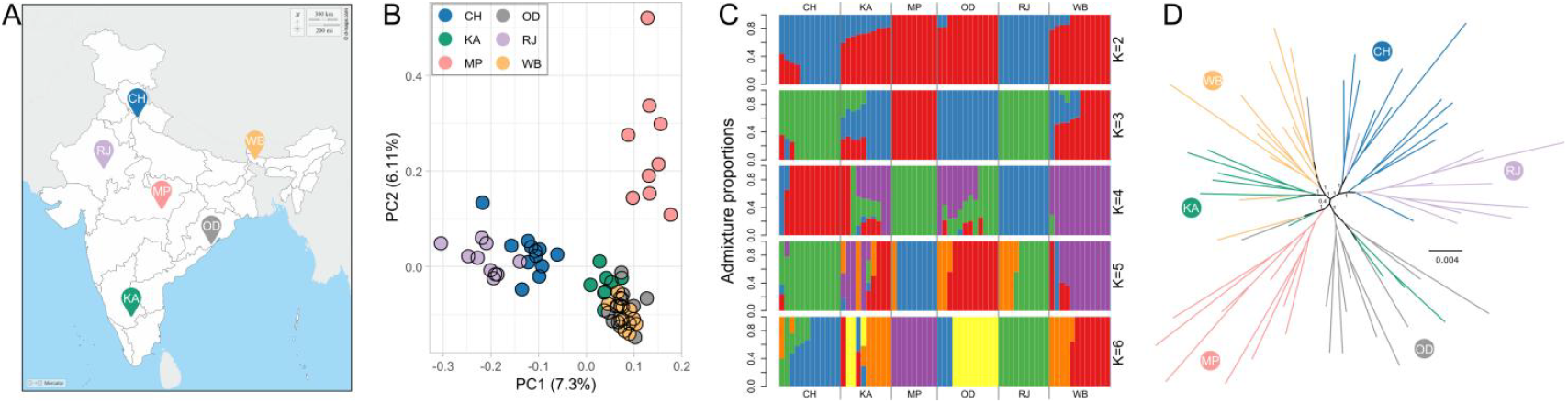
Population genetic structure inferred from genome-wide single nucleotide polymorphisms (SNPs) in parthenium beetle. (A) Approximate geographic locations of sampled populations across India. Sampling sites are labeled using region codes (see Table S1). The map is for illustrative purposes only. It was prepared using the free base map retrieved from https://www.d-maps.com/carte.php?num_car=4182&lang=en#google_vignette on 24 Apr 2026. (B) Principal component analysis (PCA) of 65 individuals from six regions, showing clustering along the first two principal components (PC1 and PC2). (C) Admixture analysis depicting individual ancestry proportions for K = 2–6. (D) Approximate maximum-likelihood phylogenetic tree inferred using FastTree from genome-wide SNP data. Node labels indicate Shimodaira–Hasegawa (SH)-like local support values. Branches are colored as per the sampling regions.

### Genetic structure

Principal component analysis (PCA) based on 4,67,071 linkage disequilibrium (LD)-pruned SNPs revealed signatures of genetic structuring within the dataset (Fig. 1B; Fig. S1). However, differentiation among clusters was generally subtle, with individuals distributed along a continuum of variation while still showing partial separation from adjacent groups. Along the first principal component (PC1), individuals from RJ occupied one extreme of the continuum and showed moderate overlap with individuals from CH. The remaining samples were further separated along PC2 into two groups: one comprising individuals from MP and the other including individuals from KA, OD, and WB. Population structure was further examined using ADMIXTURE, with the number of ancestral populations (K) ranging from 2 to 6 (Fig. 1C). At K = 2, individuals from RJ and MP formed distinct ancestry components, whereas individuals from all other regions exhibited varying degrees of admixture. At K = 3, OD was recovered as an additional ancestral component, while individuals from WB, CH and KA largely remained admixed. Consistent with their proximity in PCA space, CH individuals showed a higher proportion of the RJ ancestry component, whereas KA exhibited the highest degree of admixture across multiple lineages. At higher values (K = 4 to K = 6), admixed individuals were observed across nearly all regions, except RJ and MP at K = 4 and 6. Cross-validation indicated K = 1 as the optimal model; however, Q-statistics comparison using FSTruct suggested that K= 6 provided a better fit (p < 2e-16) (Fig. S2).

Overall, these results indicate weak to moderate genetic structuring across populations. Phylogenetic relationships inferred using FastTree2, based on concatenated LD-prunned SNPs, were broadly consistent with these patterns (Fig. 1D). Individuals from MP and RJ (with the exception of one RJ sample) formed monophyletic clusters corresponding to their geographic origins. CH individuals were placed close to RJ in correspondence to their overlapping in the PCA graph. Similarly, few individuals from KA were distributed across WB and OD groups, supporting the PCA-based clustering pattern.

### Genetic diversity

We quantified genetic diversity across the six regions using nucleotide diversity (π) and heterozygosity (H) (Fig. S3). Nucleotide diversity was estimated at the regional level, whereas heterozygosity was calculated for each individual. Differences in these metrics among regions were assessed using the Wilcoxon rank-sum test. Nucleotide diversity varied across regions, with the highest mean value observed in MP and CH (π = 0.0012) and the lowest in RJ (π = 0.001) (Table 1). Pairwise comparisons indicated significant differences in nucleotide diversity among all regions (Wilcoxon rank-sum test, p < 0.05), except between KA and OD (p=0.79). In contrast, average heterozygosity across genome did not differ significantly among regions (Wilcoxon rank-sum test, p > 0.05), with regional mean values ranging from 0.0006 to 0.0007 (Table 1). Tajima’s D values were negative across all regions, ranging from -1.09 in KA to -0.85 in MP (Table 1; Fig. S3), consistent with an excess of low-frequency variants. To assess population differentiation, we estimated pairwise genetic differentiation (fixation index, F_ST_) and absolute genomic divergence (d_XY_). Genome-wide mean pairwise F_ST_ values ranged from 0.005 (between KA and OD, KA and WB) to 0.026 (between RJ and MP), indicating low overall differentiation (Fig. S4). In contrast, mean d_XY_ values were relatively uniform across all pairwise comparisons, ranging from 0.134 to 0.143 (Fig. S4).

**Table 1.**
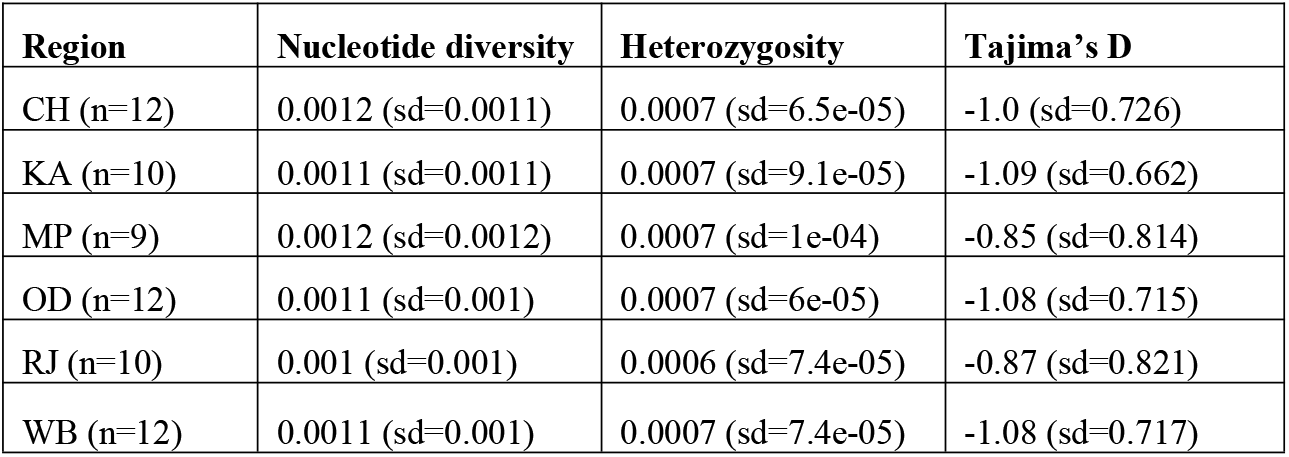
Summary statistics of genetic diversity across six regions. Mean nucleotide diversity, observed heterozygosity, and Tajima’s D are reported for each region. ‘n’ denotes the number of individuals, and ‘sd’ indicates the standard deviation.

### Inbreeding pattern

Analysis of runs of homozygosity (ROH) in the dataset did not detect long ROH segments: the maximum observed ROH length was 475 kb; yet, 91-98% of all ROHs in an individual remained within 100 kb length (Fig. 2A). Across the 25–475 kb range, MP exhibited the highest number of ROHs and cumulative ROH proportion (F_ROH_), with an average of 1212 segments spanning 13% of the genome (Fig. 2B; S5). In contrast, KA showed the lowest values, with a mean of 685 ROHs and an F_ROH_ of 6.8% (Fig. 2B; S5). However, F_ROH_ differences among all six regions were not statistically significant (Wilcoxon rank-sum test, p > 0.05). Analyses of ROHs > 100 kb showed that MP exhibited the highest F_ROH_, with an average of 2.13% of the genome, whereas KA exhibited the lowest F_ROH_, with only 0.57% of the genome in this ROH class (Fig. 2C).

**Figure 2.**
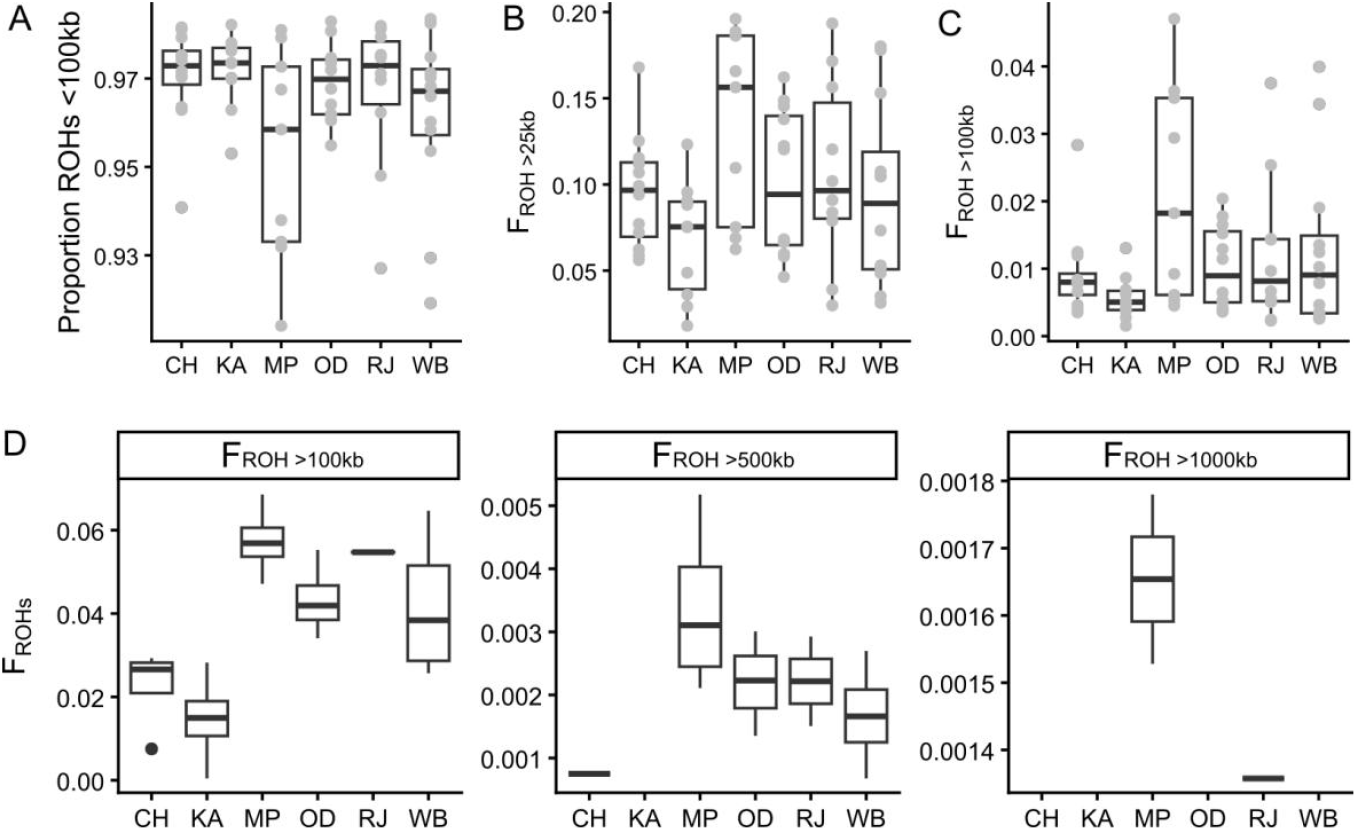
Summary of runs of homozygosity (ROH >25 kb) across six regions. (A) Box plot showing the proportion of short ROHs (<100 kb) relative to the total count of ROHs (>25 kb). (B) Box plot of cumulative ROH (F_ROH_) per individual across regions. (C) Box plot of F_ROH_ >100 kb per individual across regions. (D) Region-wise F_ROH_ partitioned by ROH length classes (>100 kb, >500 kb, and >1000 kb) from the analyses of high-coverage samples.

The absence of ROHs > 475 kb in our dataset may reflect the influence of genotype uncertainty and filtering thresholds applied to the combined data. To evaluate this possibility, we reanalyzed the high-depth dataset independently, which comprised comparatively high quality genotypes. This analysis resulted in a substantial increase in the number of detected ROHs; for example, in MP, the mean number of ROHs > 25 kb increased to 2052 segments (Fig. S5). Despite this increase, the majority of ROHs (>87%) remained within the shorter length class (< 100 kb) (Fig. S5), consistent with the pattern observed in the combined dataset (Fig. 1A). These shorter ROHs likely arise from background recombination and the fragmentation of ancestral long ROHs.

To specifically assess recent inbreeding, we focused on long ROHs (> 100 kb) from high-depth dataset. As shown in Fig. 2D, MP exhibited the highest F_ROH_, with an average of 5.74% of the genome contained in ROHs > 100 kb and 0.3% in ROHs > 500 kb. RJ showed comparable values (5.47% and 0.2%, respectively), whereas KA exhibited the lowest F_ROH_, with only 1.46% of the genome in ROHs > 100 kb and no detectable ROHs > 500 kb. Very long ROHs (> 1 Mb) were rare and observed only in MP and RJ spanning 0.16% and 0.14% genome-wide, respectively. Overall, the predominance of short ROHs and the scarcity of long segments indicate limited recent inbreeding across populations. This inference is further supported by additional metrics, including individual inbreeding coefficients, which consistently indicate low levels of contemporary inbreeding across all six regions.

### Demographic history

We inferred recent demographic history using a linkage disequilibrium (LD)-based approach implemented in the software GONe (Coombs et al., 2012). The analyses revealed marked regional differences in effective population size (N_e_) trajectories over the last 400 generations (Fig. 3), particularly during the interval spanning 50–400 generations ago, corresponding to approximately 40 years before present and closely matching the documented introduction period (∼36 years ago). We interpreted estimates from the most recent interval (0–50 generations) with caution because the consistently declining N_e_ trend across regions may partly reflect fine-scale population subdivision rather than true demographic contraction (Bansal and Nichols, 2025). Such subdivision is plausible given the discontinuous distribution of beetle populations associated with the patchy occurrence of the host plant *P. hysterophorus*. Consistent with this interpretation, each regional dataset comprised samples from two localities separated by 4–40 km (Table S1), supporting the existence of within-region genetic structure.

**Figure 3.**
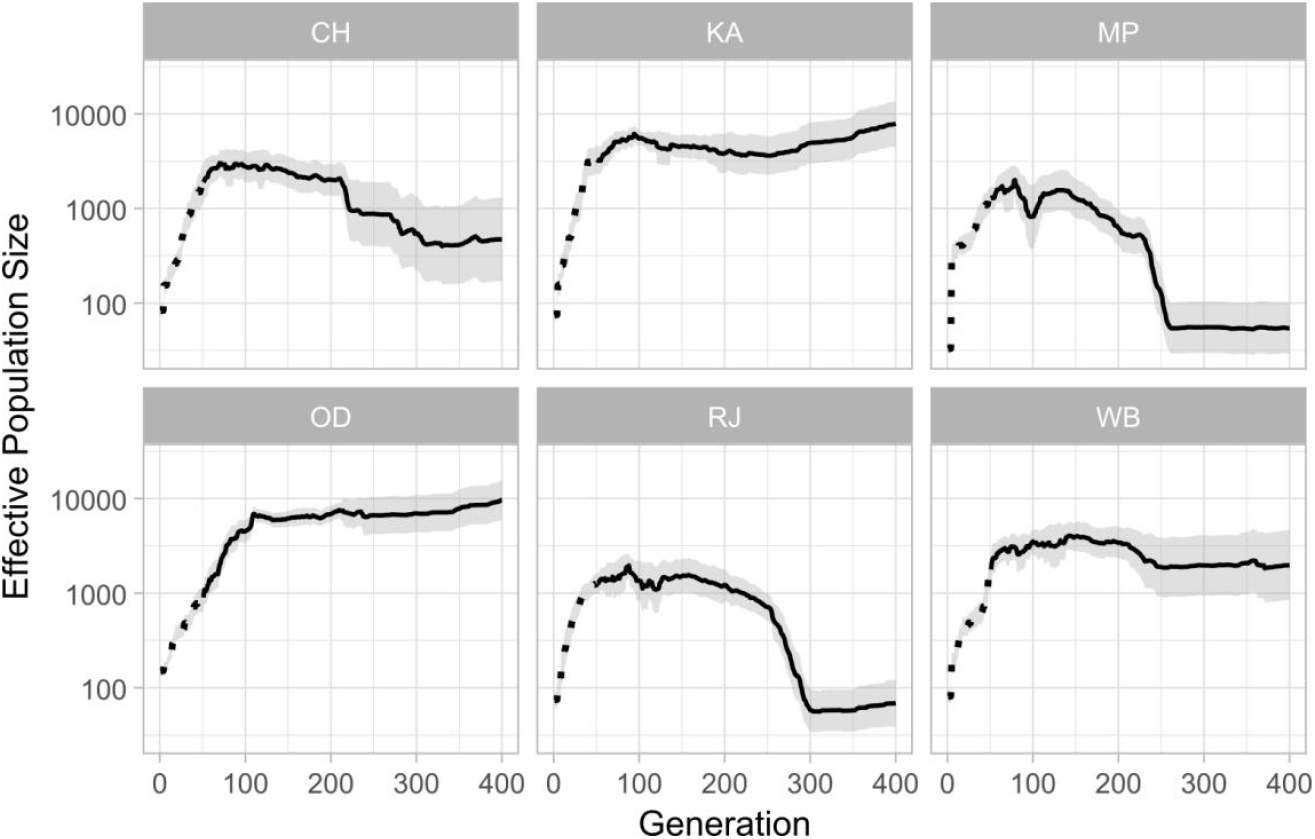
Historical changes in effective population size (N_e_) over the past 400 generations inferred using GONe. Estimations for recent 0-50 generations are marked with dotted line. Effective population size trajectories were estimated from genome-wide SNP data derived from the 187 longest contigs. Additional results are in Fig. S6.

During the 50–400 generation interval, KA, the region of initial introduction, maintained consistently high N_e_ despite the demographic bottleneck and laboratory inbreeding experienced prior to field release (Fig. 3). OD and WB, which formed a closely related genetic cluster with KA (Fig. 1B), exhibited broadly similar demographic trajectories characterized by relatively high and stable N_e_ (Fig. 3). In contrast, CH showed comparatively lower initial N_e_, followed by a gradual increase and subsequent stabilization over time. The most pronounced demographic changes were observed in MP and RJ, the two regions that formed distinct genetic clusters and exhibited the highest pairwise genetic differentiation relative to the remaining regions (Fig. 1B; Fig. S4). Both regions showed substantially reduced initial N_e_, indicative of strong founder effects or severe bottlenecks during establishment, followed by gradual demographic recovery across subsequent generations.

Apart from the relative trajectories of N_e_, the absolute magnitude of inferred effective sizes appeared substantially higher than the documented founder census sizes. For example, despite reports that the laboratory culture used for the initial release in KA was established from 66 founding individuals, the earliest inferred N_e_ in this region was in the order of 10^3^. This discrepancy may reflect the weak correspondence between census and effective population size in recently established populations. Several demographic factors could contribute to such elevated estimates, including the retention of ancestral source diversity, rapid post-introduction population expansion, or insufficient time since introduction for genetic drift to generate the strong linkage disequilibrium expected under a severe bottleneck.

## Discussion

To understand the genomic consequences of biocontrol introduction and breeding-based augmentation practices, we employed reference-based population genomics in an exemplar biocontrol system, *C. bicolorata*. This work provides a critical foundation for advancing genome-informed biocontrol programs by characterizing the current population genomic status of the species within one of its introduced ranges.

Our analyses indicate that contemporary beetle populations in India lack strong discrete population structure across sampled regions (optimal K = 1; maximum pairwise F_ST_ < 0.03; Fig. S2, S4), despite approximately 36 years since introduction (Jayanth, 1987) and extensive spread across diverse eco-climatic regions (Gharde et al., 2019). Such limited differentiation following introduction is likely attributable to a shared source population, recent common ancestry, and recurrent processes including augmentation practices and natural dispersal capacity. Nevertheless, the clustering patterns observed in PCA (Fig. 1B), together with Q-statistic comparisons (Fig. S2) and pairwise F_ST_ estimates (Fig. S4), suggest the presence of subtle population structure, particularly involving the MP and RJ populations. These patterns indicate slight shifts in allele composition among regions rather than strong population subdivision. As discussed below, elevated levels of inbreeding in these regions may have contributed to their modest genetic differentiation.

Beyond patterns of population structure, analysis of genomic variation revealed that beetle populations across India maintain relatively low levels of genetic diversity compared to other insect biocontrol agents and invasive species. In the absence of genomic data from historical samples or source populations, we contextualized the observed diversity by comparing it with insect species for which population-level genomic data are available. This comparison reveals that the highest region-wise heterozygosity in *C. bicolorata* is approximately 7.1-fold lower than the lowest reported values in populations of Colorado potato beetle *Leptinotarsa decemlineata* (a pest of *Solanum tuberosum*) in the United States (H = 0.005–0.007) (Crossley et al., 2019). Likewise, populations of *Ostrinia furnacalis* from China exhibit even higher heterozygosity, ranging from 0.0079 to 0.019 (Peng et al., 2023). Consistent with these patterns, region-wise nucleotide diversity in *C. bicolorata* is at least 5-fold lower than that observed in *L. decemlineata* populations (π = 0.006–0.008) (Crossley et al., 2019) and up to 27.9-fold lower than in introduced populations of tamarisk beetles (*Diorhabda* spp.) in North America (π = 0.0335) (Stahlke et al., 2021).

The relatively reduced genetic diversity in *C. bicolorata* across its introduced range in India, together with its origin from a pronounced demographic bottleneck (reportedly 66 founding individuals) (Sushilkumar, 2009), suggests that contemporary populations are characterized by lower site-wise allele frequencies and a reduced number of segregating sites. Such a genomic signature is consistent with expectations under strong founder effects and may indicate constrained evolutionary potential, particularly in the context of environmental adaptation (Sentis et al., 2022). However, such inference from inter-species comparisons should be interpreted with caution, as species-specific demographic histories and distinct evolutionary trajectories can substantially shape patterns of genetic diversity. Assessing whether this system represents a “true paradox” of invasion requires benchmarking contemporary variation against that of the source or founding population. Such comparisons would clarify whether the observed diversity reflects a substantial loss relative to the ancestral state or simply mirrors low initial standing variation. For example, the magnitude of standing genetic variation at the time of introduction may have been a primary determinant of the present-day diversity landscape. In addition, genetic purging under repeated inbreeding during mass-rearing phases could have further reduced deleterious variation while also influencing overall diversity levels.

Conversely, post-introduction population expansion and the accumulation of new mutations may have partially replenished genetic variation, offsetting losses incurred during bottlenecks and artificial propagation (Ahrens et al., 2026). Consistent with this expectation, we detect signatures of temporal changes in effective population size. Following the initial introduction – marked by a reduced census size – the population appears to have undergone rapid expansion, eventually attaining a relatively stable size over subsequent generations. However, regional differences in N_e_ trajectories suggest that local constraints have shaped demographic outcomes. A particularly notable pattern is the reduced initial N_e_ and delayed population recovery observed in MP, the region where an intensive breeding program was initiated around 2001 (∼230 generations ago) to augment field populations and accelerate spread (Sushilkumar, 2009). In line with this intervention, the MP population also exhibits a higher proportion of genome-wide runs of homozygosity (ROH), indicative of elevated inbreeding. Together, these observations suggest that while rapid increases in census size can facilitate the accumulation of new variation, the prevalence of inbred individuals critically modulates the relationship between census size and effective population size. Specifically, repeated augmentation with inbred individuals may inflate census size over short timescales without proportionally increasing effective population size. Nonetheless, this elevated census size can provide a larger mutational input in subsequent generations, potentially enabling a delayed but rapid increase in effective size. We propose that this demographic dynamic underlies the broadly consistent pattern of increasing N_e_ across regions, despite variation in the timing and magnitude of recovery.

## Methods

### Sampling and sequencing

We sampled 74 individuals of *Calligrapha* (*Zygogramma*) *bicolorata* during 2021–2022 from six regions across its distribution in India. The regions were designated based on state or city names for ease of reference: Chandigarh and Kalka (CH), Karnataka (KA), Madhya Pradesh (MP), Rajasthan (RJ), West Bengal (WB), and Odisha (OD) (see Table S1 for details). Within each region, two locations separated by a minimum distance of 4 km were sampled to capture local variability. Samples were surface-sterilized using 95% isopropanol prior to DNA extraction. High molecular weight genomic DNA was extracted from thoracic tissue following the protocol described in Sahoo et al. (2023). For a subset of samples, thoracic tissue was supplemented with head and leg tissues to ensure sufficient DNA yield. PCR-free DNA libraries with an average insert size of ∼450 bp were constructed and sequenced on the Illumina NovaSeq 6000 platform (Next Generation Sequencing facility at CSIR–CCMB, Hyderabad, India), generating paired-end reads of 150 bp. A total of 24 samples (four per region) were sequenced at high coverage with an average depth of ∼30X (high-coverage dataset), while the remaining 50 samples were sequenced at approximately 6X coverage (medium-coverage dataset). High-coverage dataset was downsampled to 6X for joint variant call with the medium-coverage dataset. Raw sequencing data generated in this study are publicly available under BioProject PRJNA1010988.

### Variant calling

Raw sequencing reads were quality trimmed using fastp v0.22 (Chen et al., 2018), retaining reads with a minimum length of 50 bp. Quality-controlled reads were aligned to the *C. bicolorata* nuclear reference genome (GCA_032362365; Sahoo et al., 2023) using BWA v0.7.18 (Li and Durbin, 2009). The resulting alignments were processed using SAMtools v1.20 (Danecek et al., 2021) to sort BAM files, remove duplicate reads, and generate index files. Variant calling was performed separately for the deep-coverage and low-coverage datasets using BCFtools v1.20 (Danecek et al., 2021), employing the ‘mpileup’ and ‘call’ commands with default parameters, and retaining variant sites only. Initial variant filtering was conducted using VCFtools v0.1.16 (Danecek et al., 2011) with the following criteria: retention of only biallelic single nucleotide variants (SNVs), minimum site quality (QUAL) > 20, minimum genotype quality (GQ) > 10, maximum missingness across individuals < 8%, minor allele count (MAC) ≥ 3, and Hardy–Weinberg equilibrium exact test p-value > 0.05. Depth-based filtering was applied to retain variants with read depth between 3X–7X. Additionally, for specific analyses, variant calling and filtering were conducted for the original high-coverage dataset following the above criterias, except the following changes: QUAL >30, GQ >30, and read depth 5X-37X. We excluded variants located within repetitive regions and sex-linked scaffolds. Repetitive regions were previously annotated for the reference genome (Sahoo et al., 2023). Sex-linked scaffolds were inferred using read-depth differences from the deep-coverage dataset via a Bayesian approach implemented in BeXY v1.0 (Caduff et al., 2024). Further filtering was performed using BCFtools to remove variants located within predicted repeat regions, sex-linked scaffolds, and contigs shorter than 100 kb. Finally, relatedness among individuals was assessed using PLINK v2.0 (Chang et al., 2015), and nine individuals exhibiting relatedness up to second-degree relationships (kinship coefficient>0.088) were excluded from downstream analyses.

### Population structure

To minimize the effects of linkage disequilibrium (LD), SNVs were pruned across all individuals using PLINK v1.9 (Chang et al., 2015) with the ‘--indep-pairwise’ function, applying a window size of 100 variants, a step size of 10 variants, and an LD threshold (r^2^) of 0.5. Principal component analysis (PCA) was conducted using PLINK v1.9 on the LD-pruned dataset to assess genetic clustering among individuals. Population structure was further inferred using ADMIXTURE v1.3 (Alexander et al., 2009). We evaluated ancestral components for K values ranging from 1 to 6, performing cross-validation to identify the optimal K. Furthermore, Q-matrices were analyzed using R package FSTruct v1.0 to compare ancestry variation across K values (Morrison et al., 2022). For phylogenetic reconstruction, LD-pruned SNVs were converted from VCF to FASTA format using the ‘vcf2phylip.py’ script (Ortiz, 2019). A maximum likelihood (ML) phylogenetic tree was reconstructed using FastTree v2.1.11 (Price et al., 2010) under the general time-reversible (GTR) model of nucleotide substitution.

### Diversity and divergence

From the final merged VCF file, region-wise variant datasets were generated using BCFtools v1.13. Each regional dataset was analyzed independently to estimate nucleotide diversity (π), heterozygosity, and Tajima’s D using VCFtools v0.1.16 (Danecek et al., 2011) with default parameters. Nucleotide diversity (π) and Tajima’s D were calculated using a sliding window approach with a window size of 10 kb. Individual genome-wide heterozygosity was estimated as the proportion of heterozygous sites relative to the total number of variant sites per individual. Differences in diversity measures among regions were assessed using the Wilcoxon rank-sum test. Population genetic differentiation (F_ST_) between all pairwise combinations of populations was calculated using VCFtools based on a 10 kb sliding window framework. Absolute genomic divergence (d_XY_) between population pairs was estimated using the ‘popgenWindows.py’ script (github.com/simonhmartin/genomics_general), employing non-overlapping sliding windows of 50 kb and requiring a minimum of 50 variant sites per window.

### Inbreeding and effective size analyses

Runs of homozygosity (ROH) were identified using the R package detectRUNS v0.9.6 (Biscarini et al., 2019). The VCF file was converted into PLINK-compatible genotype and map files using PLINK v1.9. These files were subsequently analyzed in detectRUNS to identify ROH segments longer than 25 kb. ROH detection was performed using the following parameters: a sliding window size of 50 variant sites, a minimum of 20 SNVs per ROH, and a minimum marker density of 1 SNV per kb. All other parameters were retained at their default settings. Recent demographic history in terms of effective population size (N_e_) was inferred using a linkage disequilibrium (LD)-based approach implemented in GONe programme (Coombs et al., 2012). Analyses were conducted over 400 generations. Default parameters were used except for the following: the recombination rate was set to 4.8 cM/Mb, based on estimates reported for related taxa in Curculionoidea (Nadachowska-Brzyska et al., 2026), and the ‘hc’ parameter was set to 0.01 to account for potential recent gene flow among populations.

## Author Contributions

RKS: Conceptualization, Data curation, Formal analysis, Funding acquisition, Investigation, Methodology, Project administration, Resources, Supervision, Validation, Visualization, Writing – original draft, and Writing – review and editing.

KV: Resources, and Writing – review and editing.

## Acknowledgements

This work was supported by the DST–INSPIRE Faculty Fellowship to RKS (DST/INSPIRE/04/2019/000478). RKS thanks the researchers and field staff who assisted with field sampling of the study species. We thank the high performance computing facility at CSIR–CCMB, Hyderabad.

## Data availability

Sequences generated in this study are submitted in GenBank, and are available under the BioProject PRJNA1010988 (SRR38359175–SRR38359219, SRR38400736–SRR38400739, SRR38401319– SRR38401322, SRR38407181–SRR38407188, SRR38420820–SRR38420823, SRR38424395– SRR38424398). Additional information and resources may be obtained from the lead contact (Ranjit Kumar Sahoo).

## References

1. Ahrens CW, Miller AD, Silver LW, McLennan EA, Hogg CJ, Weeks AR. 2026. Escaping bottlenecks: The demographic path to genetic recovery in koalas (Phascolarctos cinereus). Science. 391(6789):1010–1014.

2. Alexander DH, Novembre J, Lange K. 2009. Fast model-based estimation of ancestry in unrelated individuals. Genome Res. 19(9):1655–64.

3. Bansal JK, Nichols RA. 2025. Can genomic analysis actually estimate past population size? Trends Genet. 41(7):559–567.

4. Biscarini F, Cozzi P, Gaspa G, Marras G. 2019. detectRUNS: An R Package to Detect Runs of Homozygosity and Heterozygosity in Diploid Genomes. 10.32614/CRAN.package.detectRUNS.

5. Brodeur J. 2012. Host specificity in biological control: insights from opportunistic pathogens. Evol Appl. 5(5):470–80.

6. CABI. 2021. Datasheet Zygogramma bicolorata (Mexican beetle). CABI Compendium. 10.1079/cabicompendium.57506.

7. Caduff M, Eckel R, Leuenberger C, Wegmann D. 2024. Accurate Bayesian inference of sex chromosome karyotypes and sex-linked scaffolds from low-depth sequencing data. Molecular Ecology Resources, 24:e13913.

8. Chang CC, Chow CC, Tellier LC, Vattikuti S, Purcell SM, Lee JJ. 2015. Second-generation PLINK: rising to the challenge of larger and richer datasets. Gigascience. 4:7.

9. Chen S, Zhou Y, Chen Y, Gu J. 2018. Fastp: an ultra-fast all-in-one FASTQ preprocessor. Bioinformatics 34:i884–i890.

10. Cock MJ, van Lenteren JC, Brodeur J, Barratt BIP, Bigler F, et al. 2010. Do new access and benefit sharing procedures under the convention on biological diversity threaten the future of biological control? BioControl 55:199–218.

11. Coombs JA, Letcher BH, Nislow KH. 2012. GONe: software for estimating effective population size in species with generational overlap. Mol Ecol Resour. 12(1):160–3.

12. Crossley MS, Rondon SI, Schoville SD. 2019. Patterns of genetic differentiation in Colorado potato beetle correlate with contemporary, not historic, potato land cover. Evol Appl. 12(4):804–814.

13. Danecek P, Auton A, Abecasis G, Albers CA, Banks E, DePristo MA, Handsaker RE, Lunter G, Marth GT, Sherry ST, McVean G, Durbin R;1000 Genomes Project Analysis Group. 2011. The variant call format and VCFtools. Bioinformatics. 27(15):2156–8.

14. Danecek P, Bonfield JK, Liddle J, Marshall J, Ohan V, Pollard MO, Whitwham A, Keane T, McCarthy SA, Davies RM, Li H. 2021. Twelve years of SAMtools and BCFtools. Gigascience. 10(2):giab008.

15. De Clercq P, Mason PG, Babendreier D. 2011. Benefits and risks of exotic biological control agents. BioControl. 56:681–698.

16. Deplazes-Zemp A, Abiven S, Schaber P, Schaepman M, Schaepman-Strub G, Schmid B, Shimizu KK, Altermatt F. 2018. The Nagoya protocol could backfire on the global south. Nature Ecology and Evolution. 2:917–919.

17. Dhileepan K, Strathie L. 2009. Parthenium hysterophorus L. (Asteraceae). In: Muniappan R, Reddy GVP, Raman A, editors. Biological control of tropical weeds using arthropods. Cambridge: Cambridge University Press. p. 274–318.

18. Dhileepan K, Wilmot Senaratne KAD. 2009. How widespread is Parthenium hysterophorus and its biological control agent Zygogramma bicolorata in South Asia? Weed Res. 49:557–562.

19. Driesche RV, Hoddle M. 2016. Non-target effects of insect biocontrol agents and trends in host specificity since 1985. CAB Rev. 11:044.

20. Estoup A, Ravigné V, Hufbauer R, Vitalis R, Gautier M, Facon B. 2016. Is There a Genetic Paradox of Biological Invasion?. Annual Review Ecology, Evolution, and Systematics. 47:51–72.

21. Fagan, W.F., Lewis, M.A., Neubert, M.G. and Van Den Driessche, P. 2002. Invasion theory and biological control. Ecology Letters. 5: 148–157.

22. Fauvergue X, Vercken E, Malausa T, Hufbauer RA. 2012. The biology of small, introduced populations, with special reference to biological control. Evol Appl. 5(5):424–43.

23. Galli M, Feldmann F, Vogler UK. et al. 2024. Can biocontrol be the game-changer in integrated pest management? A review of definitions, methods and strategies. J Plant Dis Prot. 131:265–291.

24. Gharde Y, Sushilkumar Sharma AR. 2019. Exploring models to predict the establishment of the leaf-feeding beetle Zygogramma bicolorata (Coleoptera: Chrysomelidae) for the management of Parthenium hysterophorus (Asteraceae: Heliantheae) in India. Crop Protection. 122:57–62.

25. He M, et al. 2021. Herbivory of a biocontrol agent on a native plant causes an indirect trait-mediated non-target effect on a native insect. J Ecol. 109:2692–2704.

26. Heimpel GE, Cock MJW. 2018. Shifting paradigms in the history of classical biological control. BioControl. 63:27–37.

27. Hoelmer KA, and Kirk AA. 2005. Selecting arthropod biological control agents against arthropod pests: can the science be improved to decrease the risk of releasing ineffective agents? Biological Control 34: 255–264.

28. Hufbauer RA, Roderick GK. 2005. Microevolution in biological control: Mechanisms, patterns, and processes. Biological Control. 35(3):227–239.

29. Jayanth KP, Nagarkatti S. 1987. Investigations on the host specificity and damage potential of Zygogramma bicolorata Pallister (Coleoptera: Chrysomelidae) introduced into India for the biological control of Parthenium hysterophorus. Entomon. 12(2):141–145.

30. Jayanth KP. 1987. Introduction and establishment of Zygogramma bicolorata on Parthenium hysterophorus at Bangalore, India.Current Science. 56: 310–311.

31. Jayanth KP. 1991. Bio-Ecological And Physiological Studies On The Insect Zygogramma Bicolorata P And Evaluation Of Its Potential In Controlling The Weed Parthenium Hysterophorus L. PhD Thesis. Bangalore University. http://hdl.handle.net/10603/85395.

32. Li H. and Durbin R. (2009) Fast and accurate short read alignment with Burrows-Wheeler transform. Bioinformatics, 25, 1754–1760.

33. Louda SM, Kendall D, Connor J, Simberloff D. 1997. Ecological effects of an insect introduced for the biological control of weeds. Science. 277:1088–1090.

34. Mishra B. 2012. Studies on biological and reproductive behaviour of Zygogramma bicolorata pallister. Deen Dayal Upadhyaya Gorakhpur University. PhD Thesis. http://hdl.handle.net/10603/228656.

35. Morrison ML, Alcala N, Rosenberg NA. 2022. FSTruct: An FST-based tool for measuring ancestry variation in inference of population structure. Mol. Ecol. Resources. 22(7):2614–2626.

36. Nadachowska-Brzyska K, Maryańska-Nadachowska A, Kandasamy D, Andersson MN, Nowak Z, Zieliński P, Rodriguez M, Babik W. 2026. Recombination landscape shaped by inversion polymorphisms: a high-density linkage map and chromosome-level assembly of inversion-rich spruce bark beetle genome. G3 Genes|Genomes|Genetics. 16(4):jkag017.

37. Ortiz EM. 2019. vcf2phylip v2.0: convert a VCF matrix into several matrix formats for phylogenetic analysis. DOI:10.5281/zenodo.2540861.

38. Peng Y, Jin M, Li Z, Li H, Zhang L, Yu S, Zhang Z, Fan R, Liu J, Xu Q, Wilson K, Xiao Y. 2023. Population Genomics Provide Insights into the Evolution and Adaptation of the Asia Corn Borer. Mol Biol Evol. 40(5):msad112.

39. Price MN, Dehal PS, Arkin AP. 2010. FastTree 2 -Approximately Maximum-Likelihood Trees for Large Alignments. PLoS ONE. 5(3):e9490.

40. Sahoo RK, Manu S, Chandrakumaran NK, Vasudevan K. 2023. Nuclear and Mitochondrial Genome Assemblies of the Beetle, Zygogramma bicolorata, a Globally Important Biocontrol Agent of Invasive Weed Parthenium hysterophorus. Genome Biol Evol. 15(10):evad188.

41. Schaffner U, Steinbach S, Sun Y, Skjøth CA, de Weger LA, Lommen ST, Augustinus BA, Bonini M, Karrer G, Šikoparija B, Thibaudon M, Müller-Schärer H. 2020. Biological weed control to relieve millions from Ambrosia allergies in Europe. Nat Commun. 11(1):1745.

42. Schrieber K, Lachmuth S. 2017. The Genetic Paradox of Invasions revisited: the potential role of inbreeding x environment interactions in invasion success. Biol Rev Camb Philos Soc. 92(2):939–952.

43. Schwarzländer M, Hinz HL, Winston RL, et al. 2018. Biological control of weeds: an analysis of introductions, rates of establishment and estimates of success, worldwide. BioControl. 63:319–331.

44. Seastedt TR. 2015. Biological control of invasive plant species: a reassessment for the Anthropocene. New Phytol. 205(2):490–502.

45. Sentis A, Hemptinne JL, Magro A, Outreman Y. 2022. Biological control needs evolutionary perspectives of ecological interactions. Evol Appl. 15(10):1537–1554.

46. Shawn M. Clark, Hume B. Douglas, and Daniel J. Cavan. 2024. Notes on Calligrapha Chevrolat (Subgenus Zygogramma Chevrolat) and Tritaenia Motschulsky (Coleoptera: Chrysomelidae: Chrysomelinae). The Coleopterists Bulletin 78(2):281–295. doi:10.1649/0010-065X-78.2.281.

47. Stahlke AR, Bitume EV, Özsoy ZA, Bean DW, Veillet A, Clark MI, Clark EI, Moran P, Hufbauer RA, Hohenlohe PA. 2021. Hybridization and range expansion in tamarisk beetles (Diorhabda spp.) introduced to North America for classical biological control. Evol Appl. 15(1):60–77.

48. Stenberg JA, Sundh I, Becher PG. et al. 2021. When is it biological control? A framework of definitions, mechanisms, and classifications. J Pest Sci. 94:665–676.

49. Sushilkumar. 2009. Biological control of Parthenium in India: status and prospects. Indian Journal of Weed Science. 41(1&2):1–18.

50. Thomas MB, Reid AM. 2007. Are exotic natural enemies an effective way of controlling invasive plants? Trends Ecol Evol. 22(9):447–53.

51. Thomas MB, Willis AJ. 1998. Biocontrol—risky but necessary? Trends Ecol Evol. 13:325–329.

52. van Wilgen BW, Raghu S, Sheppard AW, Schaffner U. 2020. Quantifying the social and economic benefits of the biological control of invasive alien plants in natural ecosystems. Curr Opin Insect Sci. 38:1–5.

