## Supplementary material for "Population genomics of a biocontrol agent: Insights into post-introduction establishment in parthenium beetle": Tables S1 and Figures S1-S6

**Supplemental file** of the title "Population genomics of a biocontrol agent: Insights into post-introduction establishment in parthenium beetle".

**Table S1.** Details of sampling sites.

| **Locality-1** | **Locality-2** | **Approx. distance between localities (km)** | **State** | **Region code** | **Number of samples sequenced** | **Number of samples analyzed** |
|---|---|---|---|---|---|---|
| Bangalore-1 (13.112179 N 77.528635 E) | Bangalore-2 (12.845770 N 77.774524 E) | 39.84 | Karnataka | KA | 12 | 10 |
| Chandigarh-1 (30.747341 N 76.742141 E) | Kalka-1 (30.835174 N 76.964965 E) | 23.42 | Himachal Pradesh | CH | 12 | 12 |
| Ajmer-1 (26.485372 N 74.606717 E) | Ajmer-2 (26.451314 N 74.589675 E) | 4.15 | Rajasthan | RJ | 14 | 10 |
| Jabalpur-1 (23.219458 N 79.974184 E) | Jabalpur-3 (23.131040 N 79.800404 E) | 20.30 | Madhya Pradesh | MP | 12 | 9 |
| Siliguri-1 (26.743673 N 88.431149 E) | Siliguri-3 (26.684597 N 88.443433 E) | 6.68 | West Bengal | WB | 12 | 12 |
| Berhampur-1 (19.313170 N 84.827125 E) | Berhampur-2 (19.285999 N 84.786819 E) | 5.20 | Odisha | OD | 12 | 12 |

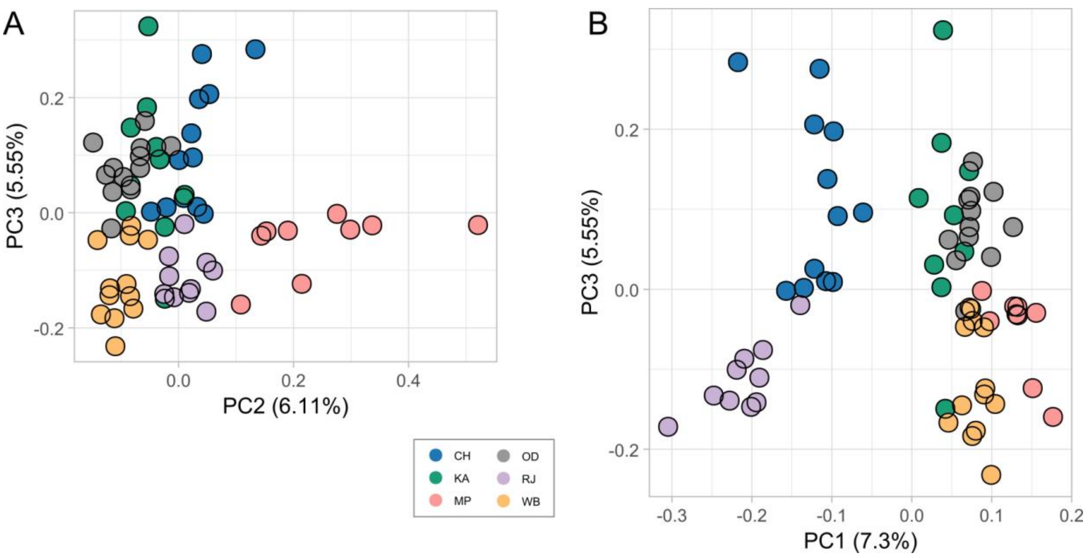


**Figure S1.** Principal component analysis (PCA) of 65 individuals from six regions along PC2–PC3 (A) and PC1–PC3 (B).

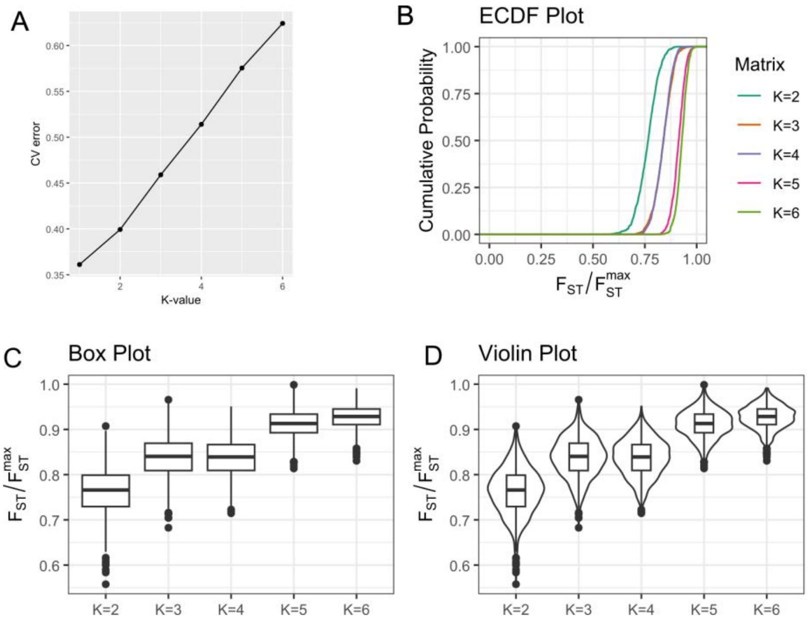


**Figure S2.** Cross-validation error and distribution of admixture proportions across K = 1–6. (A) Line plot showing cross-validation (CV) error across K values. Empirical cumulative distribution function (ECDF) curves (B), box plots (C), violin plots (D) summarize the distribution of ancestry coefficients (Q) across K. Pairwise comparisons using Wilcoxon rank sum test with continuity correction showed that, given the underlying matrix, the $F_{ST}$ ratios were significantly different for all K values ($p < 2e\text{-}16$) except between K = 3 and K = 4 ($p = 0.69$).

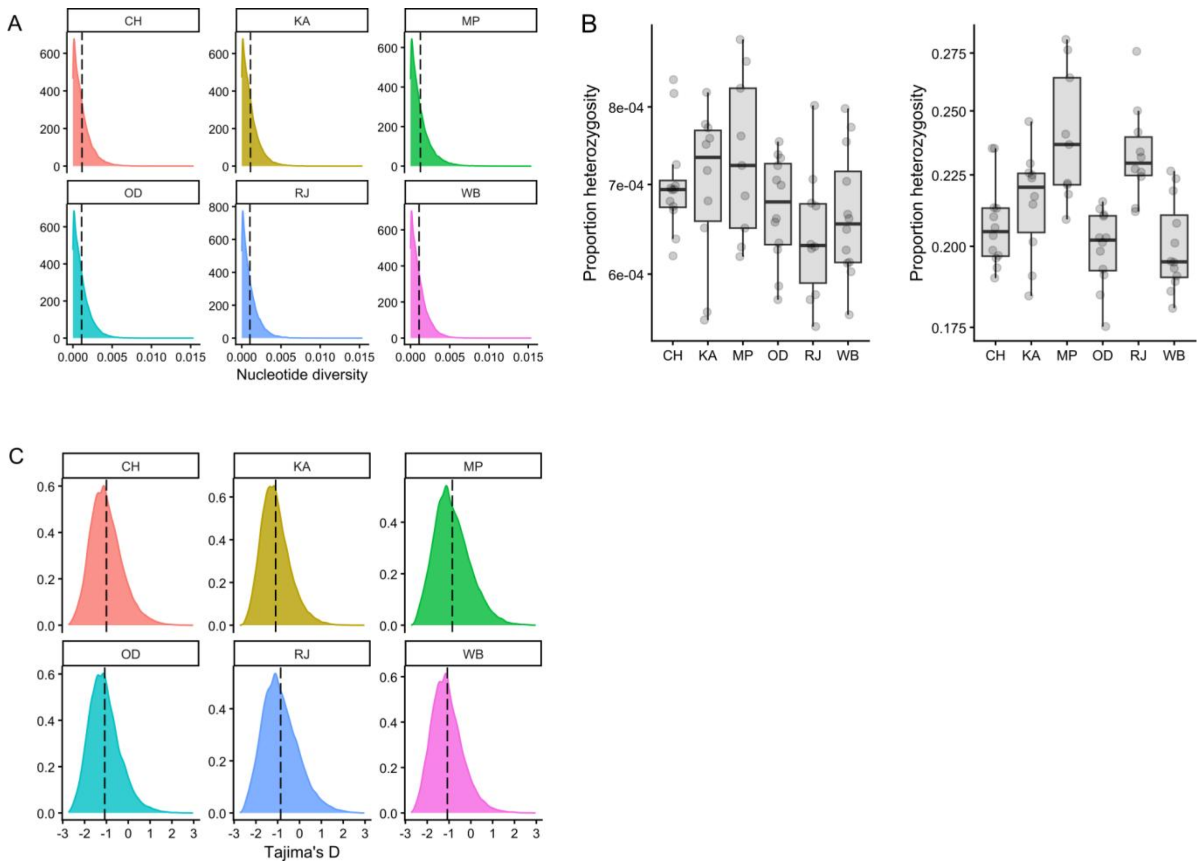


**Figure S3.** Distribution of nucleotide diversity, heterozygosity, and Tajima's D across regions. (A) Density plots showing region-wise distributions of nucleotide diversity (π), with dashed vertical lines indicating medians. (B) Bar plots showing heterozygosity (H) estimated genome-wide (left) and from variant sites only (right). (C) Density plots showing region-wise distributions of Tajima's D, with dashed vertical lines indicating medians.

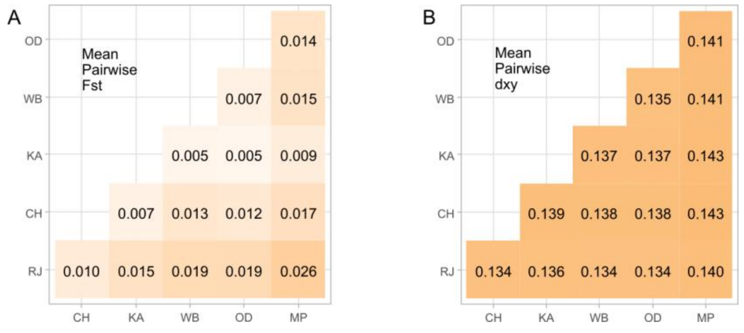


**Figure S4.** Mean pairwise genetic differentiation ($F_{ST}$) and divergence ($d_{XY}$) across regions depicted in (A) and (B), respectively.

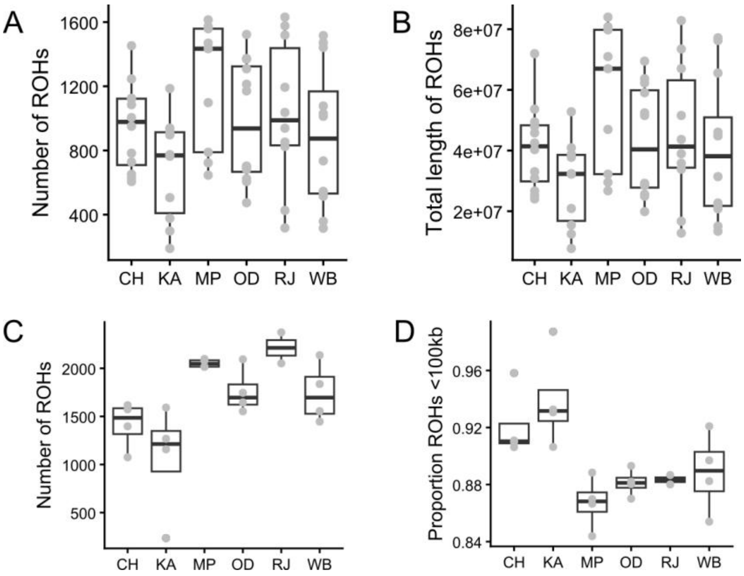


**Figure S5.** Top panel: Box plots showing the total number of ROHs (>25 kb) (A) and cumulative length of ROHs (>25 kb) (B) per individual across regions from the analyses of combined dataset. Bottom panel: Box plots showing the total number of ROHs (>25 kb) (C) and the proportion of short ROHs (<100 kb) relative to the total count of ROHs (>25 kb) (D) from the analyses of high-coverage dataset.

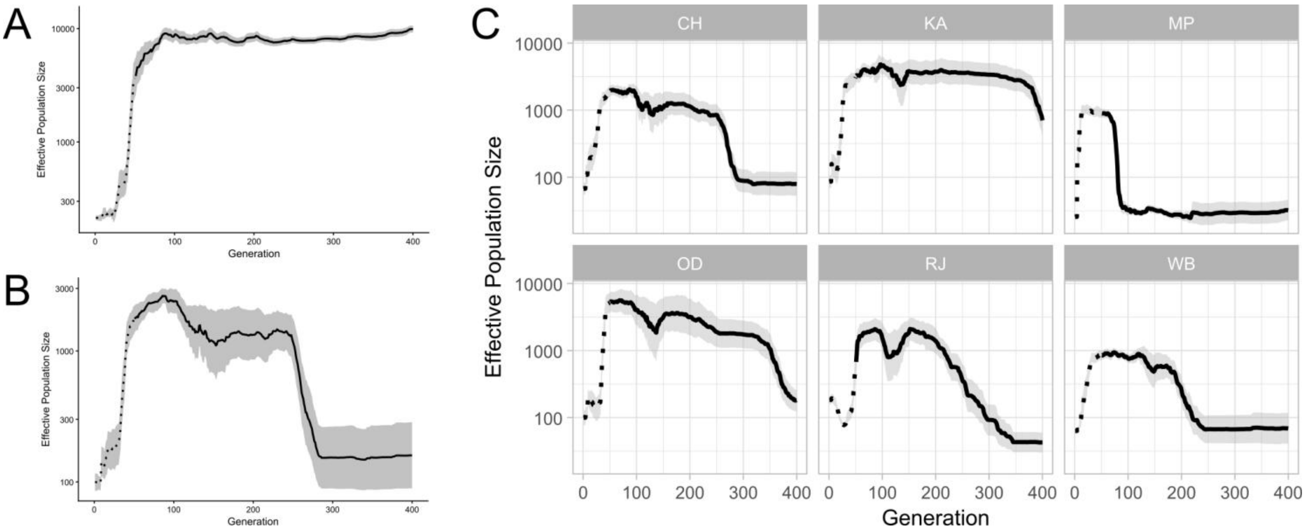


**Figure S6.** Historical changes in effective population size ($N_e$) over the past 400 generations inferred using GONe. Estimates are based on genome-wide SNPs from the 187 longest contigs. (A) Estimates based on samples pooled across all six regions. (B) Estimates based on samples pooled across all six regions, and analysed using the high-coverage dataset. (C) Region-wise estimates based on the high-coverage dataset.
